# The fate of artificial transgenes in *Acanthamoeba castellanii*

**DOI:** 10.1101/2024.10.29.620921

**Authors:** Morgan J. Colp, Cédric Blais, Bruce A. Curtis, John M. Archibald

**Affiliations:** Department of Biochemistry and Molecular Biology, Dalhousie University, Halifax, Nova Scotia Canada; Institute for Comparative Genomics, Dalhousie University, Halifax, Nova Scotia, Canada

## Abstract

In this study, artificial transformation experiments were performed to investigate how the *A. castellanii* genome responds to foreign DNA presented in both circular and linear plasmid form. Nanopore sequencing was used as a high throughput method to screen for transgene DNA in the resulting transformant cultures, and candidate transgene integrations were identified. Molecular biology experiments were performed to validate the sequence data and provide additional context on the fate of transgenes. A method was devised to estimate the rate of read chimerism in nanopore sequencing runs and accurately account for the effects of read chimerism in identifying putative transgene integrations. Based on the experimental data in hand, a potential mechanism for transgene maintenance in *A. castellanii* is proposed, one in which incoming foreign DNA is tandemly duplicated and telomeres are added to the ends. This nascent linear molecule is maintained as a transgene-bearing minichromosome, while also allowing for chromosomal integration.

## Introduction

The ability to express transgenes in eukaryotic cells has become a staple of the contemporary molecular genetics toolkit, facilitating a wide array of functional studies that answer genetic, biochemical, and cell biological questions^1,2^. These experiments often seek to tag, overexpress, or suppress proteins of interest and make inferences about that protein’s role in the organism’s biology. Sophisticated transformation systems and the genetic toolkits built upon them are available for most model organisms, and for most new species we wish to characterize on a molecular and cell biological level, researchers strive to develop a corresponding system for transformation and genetic manipulation^3^. However, while the introduction and expression of transgenes is ubiquitous in molecular biology, relatively few researchers seek to determine the fate of the introduced DNA within the host cell. This is not unjustified; for most experiments of this nature, the fate of the transgene is irrelevant to the research question. For the study of genome biology, though, this knowledge gap contains answers to questions relevant to how eukaryotic organisms maintain their own genomes and respond to the introduction of foreign DNA in nature.

In transformation experiments, evidence suggests that there are two main ways that transgenes are permanently maintained (ignoring transient transformation where the transgenes are expressed and then lost within a short time). The first option is integration into one or more loci on the host chromosomes, where the integrated DNA is faithfully replicated from generation to generation as a part of the genome. Integration into an organellar genome is formally possible, but in experiments where selection relies on expression from nuclear promoters, organellar genome integrants may not persist^4^. The second option for the stable maintenance of a transgene is for the cell to replicate it as an extrachromosomal element, or ‘episome’. Here the transgene undergoes the same replication and segregation processes over time as the host genome but remains physically separate from it. Evidence for both possibilities has been gleaned from transformation studies in a variety of different eukaryotic species, and the two options are not mutually exclusive, even in the same cell. For chromosomally integrated transgenes, additional questions arise, such as the molecular mechanism(s) of integration and the genomic features at the site of integration that may have influenced the frequency and/or process.

Here we sought to characterize the fate of artificially introduced transgenes in the amoebozoan protist *Acanthamoeba castellanii* using a transformation protocol developed by Bateman and colleagues^5,6^. The plasmids constructed for the development of this protocol contain a selectable marker, neo^R^, conferring neomycin and geneticin resistance, as well as a reporter gene, enhanced green fluorescent protein (EGFP), both expressed from endogenous *A. castellanii* promoters. Historically, answering such questions relied heavily on classical molecular tools such as polymerase chain reaction, Southern blotting, restriction fragment analysis, and Sanger sequencing as needed^7,8^. In this study, Oxford Nanopore long-read sequencing was used to perform a high-throughput, genome-wide search for transgenes in *A. castellanii,* with traditional molecular biological methods used for confirmation and additional characterization.

## Methods

### Culturing

All cultures of *A. castellanii* strain Neff used in this study were grown at room temperature in Neff base medium with additives (ATCC Medium 712; 0.75% yeast extract, 0.75% proteose peptone, 2 mM KH_2_PO_4_, 1 mM MgSO_4_, 1.5% glucose, 0.1 mM ferric citrate, 0.05 mM CaCl_2_, 1 µg/mL thiamine, 0.2 µg/mL D-biotin, and 1 ng/mL vitamin B_12_). Transformed culture media also contained the antibiotic G418 at a concentration of 10 µg/mL during initial culture establishment after transformation, and 50 µg/mL for full-strength selection and long-term maintenance.

### Transformation

All transformation experiments were based on the method described by Peng, Omaruddin, and Bateman^5^, and further developed by Bateman^6^. Our general adaptation of this protocol is available online at protocols.io^9^. The plasmid pGAPDH-EGFP, described by Bateman, was used for these experiments (Fig. 1).

**Figure 1.**
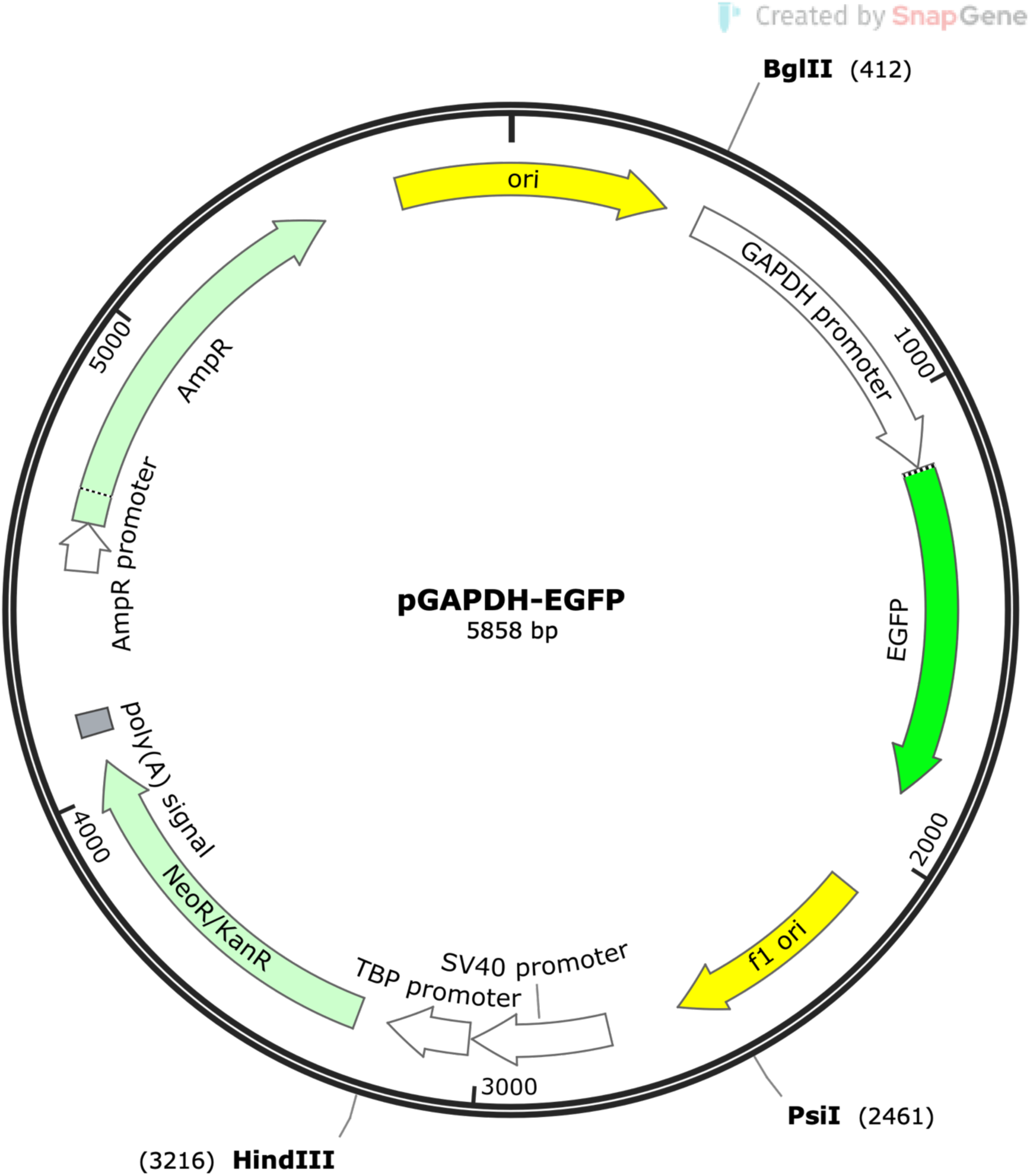
Plasmid pGAPDH-EGFP used for transformation of *Acanthamoeba castellanii*. The plasmid has an ampicillin resistance marker for propagation in *E. coli,* neomycin resistance marker for selection in *Acanthamoeba*, and enhanced green fluorescent protein (EGFP) as a reporter gene. The neomycin resistance marker and EGFP are expressed from endogenous *Acanthamoeba* promoters. Expression of the resistance marker is driven by the TATA-box binding protein (TBP) promoter, while EGFP expression is driven by the glyceraldehyde-3-phosphate dehydrogenase (GAPDH) promoter. Restriction sites used in this study are marked on the plasmid map.

### Nanopore sequencing of transformed *A. castellanii* cultures

Once the transformed cells had recovered and stable cultures were established, genomic DNA was extracted using an SDS-based lysis followed by phenol-chloroform extraction. DNA samples were cleaned with QIAGEN G/20 genomic clean-up columns using the manufacturer’s protocol, but with double the number of wash steps. A nanopore sequencing library was prepared for the Oxford Nanopore MinION using the SQK-LSK108 ligation sequencing kit and sequenced on a FLO-MIN106 flow cell. The reads were basecalled with Albacore v2.1.7.

### Bioinformatic analysis of transgene-derived DNA sequence reads

To search for reads representing potential genomic integrations of the plasmid, BLASTn v2.7.1^10^ was used to identify and retrieve all reads from the sequencing experiment with hits to the plasmid sequence. These reads were then compared against the wild-type genome assembly^11^, again using BLASTn v2.7.1 to search for plasmid-genome junctions. A simple text search was also performed against the plasmid-containing reads to search for telomeric repeats.

### Southern blot analysis to locate transgene DNA

The following samples were prepared ahead of the Southern blot experiment: undigested genomic DNA from the transformed *A. castellanii* culture, two aliquots from this gDNA that were digested with either *Bgl*II and *Hin*dIII, purified pGAPDH-EGFP plasmid, and two aliquots of the plasmid digested with the same restriction endonucleases. These samples were run on a 1% agarose gel alongside GeneRuler 1 kb plus ladder, and the DNA was transferred to a positively charged nylon membrane following the Southern blotting protocol published by ThermoFisher (https://assets.fishersci.com/TFS-Assets/LSG/manuals/MAN0013296_Southern_Blotting._Dot_Blotting_UG.pdf), which is a typical alkaline transfer method.

The probe for this experiment was generated using the Thermo Scientific Biotin DecaLabel DNA labelling kit. In this case, the probe was generated from a 345-bp segment of the neo^R^ gene on the pGAPDH-EGFP plasmid. Hybridization of this probe to the membrane also followed the ThermoFisher Southern blotting protocol. Membrane blocking was done with salmon sperm DNA at a concentration of 50 µg/mL. For the final hybridization, the probe was added to a concentration of 50 ng/mL. Hybridization was performed for 12 hours at 42 °C with gentle agitation. The membrane was then washed twice with 2X SSC + 0.1 % SDS for 10 minutes each time at room temperature and washed for high stringency in 0.1X SSC + 0.1% SDS for 10 minutes each time at 65 °C. Hybridizing probe was detected using the Thermo Scientific Biotin Chromogenic Detection kit. The detection reaction was performed according to the manufacturer’s protocol, allowing the colour to develop overnight.

### Nanopore sequencing of transformant clones

Single amoeba cells were isolated by agar plate migration and used to establish monoclonal cultures from the population of transformants^12^. Three clones were selected for nanopore sequencing, hereafter referred to as ‘Clone 1’, ‘Clone 5’, and ‘Clone 8’, based on their original labels at the time of single cell isolation. DNA was extracted from clonal cultures as described above. A barcoded sequencing library was prepared using the ligation sequencing kit (SQK-LSK109) and the native barcoding kit (EXP-NBD104) with barcodes 2, 5, and 8. The library was run on a FLO-MIN106 flow cell and basecalled with Albacore v2.3.3.

A subsequent sequencing run using only Clone 1 was sequenced with the ligation sequencing kit (SQK-LSK109) on a FLO-MIN106 flow cell. The raw output was basecalled with Guppy v3.1.5 using the HAC (high accuracy) basecalling model.

Plasmid-containing reads from each of these sequencing datasets were identified using BLASTn v2.7.1 and retrieved. They were mapped against the wild-type genome using minimap2^13^ v2.24, with soft clipping allowed. Mapped reads were then visualized in IGV to locate putative chromosomal integrations of the plasmid, and BLASTn v2.7.1 was used to confirm that the soft-clipped portion of these reads was plasmid-derived.

### Polymerase chain reaction verification of chromosomal integration of transgenes

Four putative integration loci were selected for verification with PCR, the goal being to amplify across the plasmid-genome junctions on each end of a putative integration. Early PCR experiments used NEB One *Taq* polymerase in its Quick-Load 2X master mix with standard buffer. These experiments started with the following PCR program: 96 °C for 5 minutes, then 25 cycles of denaturation at 96 °C for 30 seconds, annealing at 50 °C for 30 seconds, and extension at 72 °C for 1 minute, then a final 7-minute elongation step at 72 °C. To adjust for poor amplification, the number of cycles was then increased to 35. Then, to reduce the generation of non-specific product, the annealing temperature was increased to 56 °C.

One of the putative transgene integration loci showed promising results and PCR was further optimized using ThermoFisher Platinum II *Taq* Hot-Start DNA polymerase. First, the standard amplification recipe recommended for this polymerase was used without the optional GC enhancer reagent, which greatly improved yield. The PCR program was as follows: an initial denaturation step of 2 minutes at 94 °C, followed by 35 cycles of denaturation at 94 °C for 15 seconds, annealing at 60 °C for 15 seconds, and extension at 68 °C for 15 seconds. There was no long final extension step in this protocol. Bands of interest were gel extracted using a Macherey-Nagel NucleoSpin Gel and PCR Clean-up kit, and samples were sent to GeneWiz, South Plainfield, New Jersey for Sanger sequencing.

### Transformation experiments with linearized plasmid

Another transformation experiment was performed as described above, except with plasmid linearized by a single cut with the restriction endonuclease *Psi*I, and monoclonal cultures were established. Three of these clonal isolates were sequenced and analyzed using the same library preparation and bioinformatic approach as Clones 1, 5, and 8 described above, but basecalling was done with Guppy v5.1.13 using the SUP (super accurate) model. The three clonal isolates analyzed from this transformation experiment are hereafter referred to as ‘Clone LT6’, ‘Clone LT8’, and ‘Clone LT9’.

### Determining the rate of artefactual read chimerism in transformant sequence data

The background rate of read chimerism was estimated from each sequencing run used to infer integration. This was done using the non-plasmid-containing reads, so that the calculated rate of chimerism could be applied to assess the validity of the plasmid+genome reads. Reads from an individual sequencing run were mapped against the wild-type reference genome sequence with minimap2^13^ v2.24 and the mapping was output as a paf file rather than a SAM file. All reads that had exactly two mappings to the reference and no plasmid sequence were retained for further analysis, because depending on the genomic location of these two mappings, reads with this mapping pattern could be chimeras from two different genomic loci.

A custom Perl script was used to identify putatively chimeric reads. This script assessed the reads with two mappings to the reference genome as follows: if the two mappings for a read were found to be greater than 500 bp apart on the genome, each was more than 100 bp away from the end of a scaffold, and the distance between the two mappings on the genome was at least 100 bp more than the distance between the two mappings on the read, the read was considered to be chimeric. To estimate the proportion of chimerism in each sequencing run, the number of putative chimeric reads was divided by the total number of reads fed into the chimerism identification script.

The number of putative chromosomal integrations of the plasmid that could have been artefacts due to read chimerism was estimated using this proportion. The total number of plasmid-containing reads was multiplied by the proportion of reads expected to be chimeric, and the resulting value represented the expected number of reads to artefactually appear as putative integrations due to chimerism. This number was compared to the observed number of putative integrations from the same read set. This process was done independently for the output of each sequencing experiment to account for differing rates of chimerism across different sequencing runs.

### Illumina short-read re-sequencing and analysis of *A. castellanii* clones

Genomic DNA was extracted from wild-type *A. castellanii* as well as clones LT6 and LT9 and sent to the Integrated Microbiome Resource at Dalhousie University for Illumina library preparation and sequencing. The three samples were sequenced as part of a larger multiplexed run on an Illumina NextSeq2000 instrument. 150-bp paired-end reads were generated and processed with Trimmomatic^14^ v0.36 using the following trimming parameters: HEADCROP:10, LEADING:12, SLIDINGWINDOW:4:20, MINLEN:75.

Illumina reads from Clone LT6 and Clone LT9 were used to seek confirmation of putative integrations inferred from nanopore sequence data. The Illumina reads from each respective clone were mapped against the long reads from the same clone using HISAT2^15^ v2.2.1 with default mismatch penalties, as well as with the maximum and minimum mismatch penalties decreased from 6 and 2 to 5 and 1, respectively. Mapped reads were visualized in Integrative Genomics Viewer^16,17^ to look for reads mapping across the putative plasmid-genome junctions. Illumina reads were also compared against the putative integration-supporting long reads using BLASTn v2.7.1 to see whether any Illumina reads showed BLAST hits that spanned the junctions.

### Southern blot analysis for detection of episomal transgenes

For this experiment, the same protocol and choice of probe were used as above, but the input DNA and gel were different. Genomic DNA from wild-type *Acanthamoeba* and from Clone LT9 were used, as well as purified pGAPDH-EGFP plasmid. For each sample, one aliquot was left untreated, while a second aliquot was digested with the restriction endonucleases *Nhe*I, *Not*I, and *Sac*I. The samples were run on a 0.5% agarose gel to better resolve large DNA fragments along with GeneRuler 1 Kb ladder. The DNA was visualized with ethidium bromide, and a Southern blot experiment was performed using the same protocol as described above.

A pulsed-field gel electrophoresis (PFGE) experiment was performed on wild-type and Clone LT9 genomic DNA, with one ladder of lambda concatemers (48.5 Kbp to 1,000 Kbp) as well as the GeneRuler 1 Kb ladder. One lane contained both ladder types. The PFGE run used a 1% agarose gel, 0.5X TBE running buffer, an 18-hour run time, a 120 ° angle, and a switch time ramping from 1 second to 14 seconds over the course of the run. The gel was stained with ethidium bromide to visualize the DNA.

## Results

### Stable transformation of *Acanthamoeba castellanii*

After recovery from transformation, the transformed cell population under selection with 50 µg/mL G418 exhibited a growth rate comparable to that of a wild-type culture with no selection. Upon examination with epifluorescence microscopy, cells expressing EGFP were clearly visible (Figure 2). Successful transformation was obtained using two different plasmids that differ only in the EGFP promoter; pGAPDH-EGFP, shown in Figure 2D-E, was used in this study. In this initial transformation experiment, the circular form of the plasmid was used. Strong nuclear localization of the fluorescent protein was apparent. It is unclear why this occurred, or whether it has any biological impact on the transformed cells.

**Figure 2.**
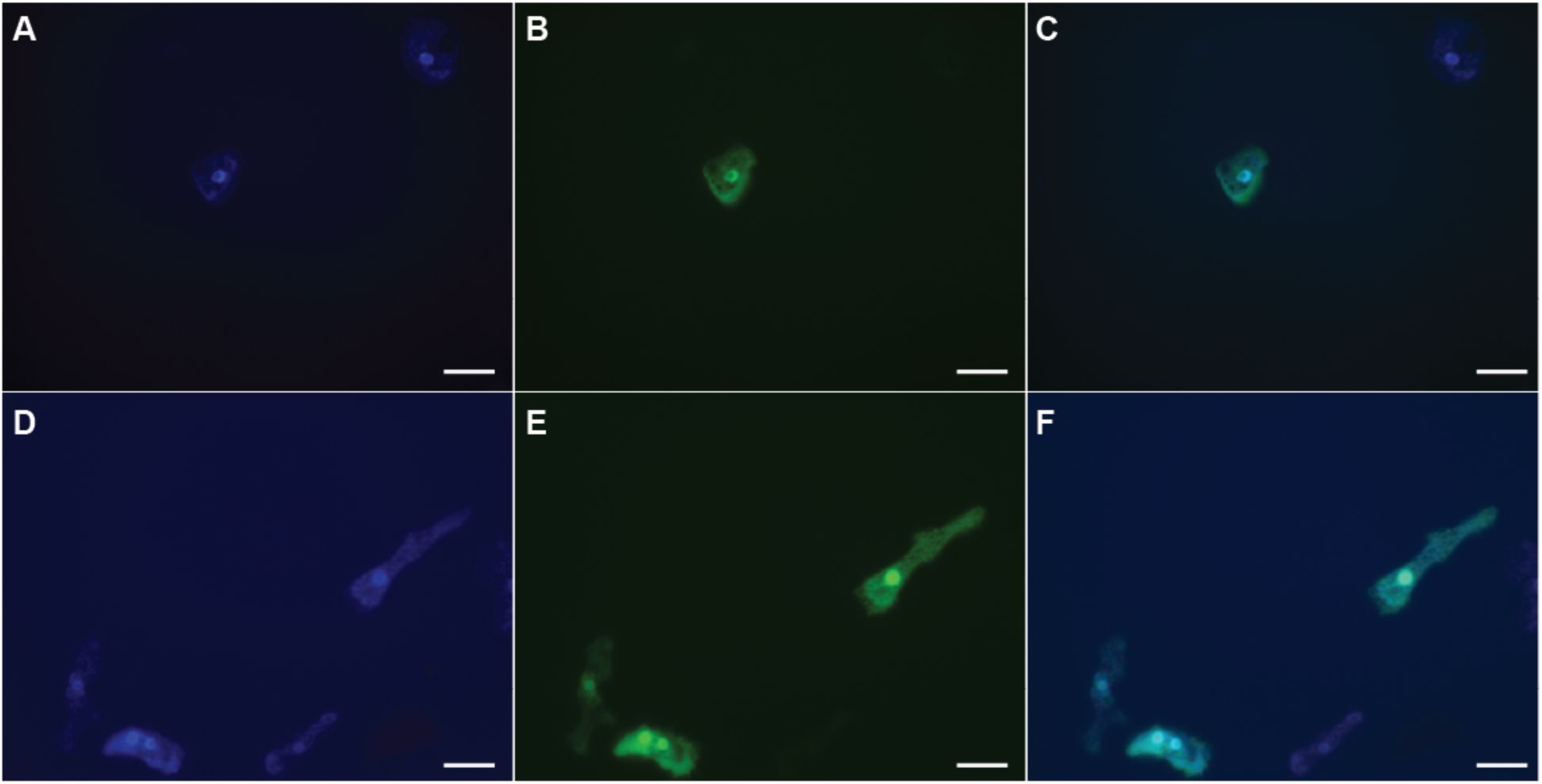
*A. castellanii* cells transfected with pTPBF-EGFP and pGAPDH-EGFP. *A-C.* Transfected with pTPBF-EGFP. *A.* Cells were imaged using fluorescence microscopy with a DAPI filter to visualize DNA stained with DAPI. *B.* Cells were imaged using fluorescence microscopy using a GFP filter to visualize EGFP expression. *C.* A merge of the DAPI and GFP images shown. *D-F.* Transfected with pGAPDH-EGFP. *D.* Cells were imaged using fluorescence microscopy with a DAPI filter to visualize DNA stained with DAPI. *E.* Cells were imaged using fluorescence microscopy using a GFP filter to visualize EGFP expression. *F.* A merge of the DAPI and GFP images shown. *A-F.* All cells were maintained under selection with 50 μg/mL G418, then fixed on slides with 4% formaldehyde, permeabilized with 0.5% Triton-X100, and stained with 300 nM DAPI for 5 minutes, then washed with phosphate-buffered saline before mounting with Fluoromount. All cells were imaged using 630X total magnification. All scale bars represent 20 μm.

### Plasmid sequence can be detected by sequencing transformants

We used Oxford Nanopore long-read sequencing as a high throughput screening tool for the detection of plasmid DNA in *A. castellanii* transformants. The sequence output corresponding to transformants stemming from transformation of circular plasmid DNA is summarized in Table 1. The goal was to look for the presence of transgene DNA in transformed cultures and to determine in what genomic context they were being maintained. BLAST searches with the pGAPDH-EGFP sequence as query allowed retrieval of all long reads corresponding to internalized plasmids. In the case of putative chromosomal integration, these were long reads containing both plasmid and genomic sequence, where at least one junction was represented. In our first experiment (‘Mixed transformants’ in Table 1), 9,579 reads were detected with identifiable plasmid sequence, of which 118 were found to have telomeric repeats on one end; a single read was found with repeats on both ends. Read mapping against the *A. castellanii* reference genome identified 33 reads that appeared to contain both genomic and plasmid sequence, and thus could correspond to chromosomal integrants. Notably, many of the plasmid-mapping reads contained tandem repeats of the single-copy plasmid sequence used in our transformation experiments, in a variety of different flanking sequence contexts. Arrays of up to 11 copies of the plasmid were observed on nanopore reads more than 65 Kbp in length; the existence of even longer arrays may have been obscured by read length limitations.

**Table 1.**
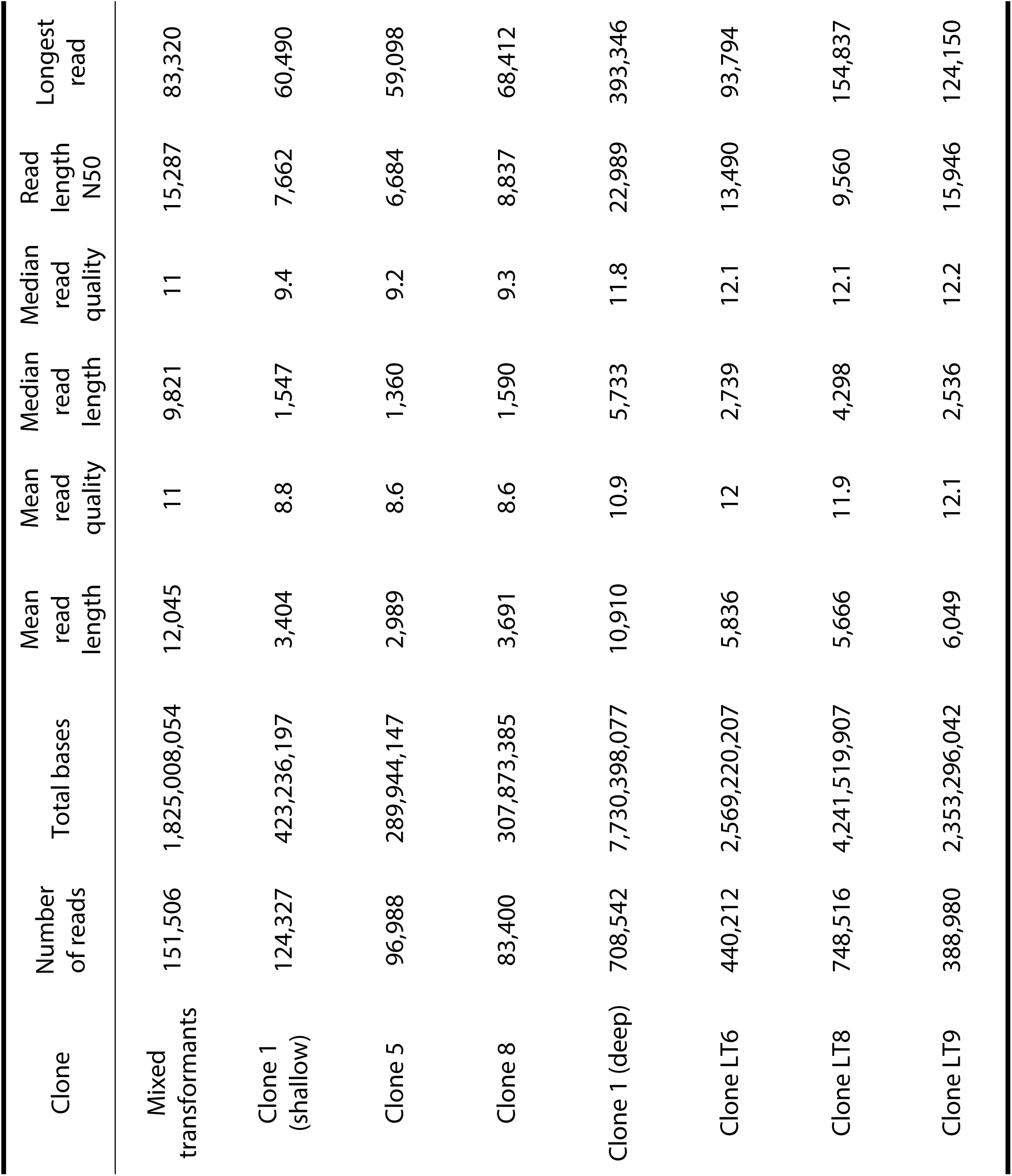
Sequencing statistics for all nanopore long-read sequencing runs performed on *Acanthamoeba castellanii* transformants in this study. The Clone 1 (shallow) read set was barcoded with Clone 5 and Clone 8 in the same run, while Clone 1 (deep) came from a separate run where Clone 1 was the only sample. Clones LT6, LT8, and LT9 were barcoded together and sequenced in one run. The mixed transformant sample was the only sample in its respective sequencing run. All the runs used FLO-MIN106 flow cells, and all used the SQK-LSK109 sequencing kit, except the mixed transformants which used the previous version, SQK-LSK108. The mixed transformants and Clones 1, 5, and 8 were transformed with circular plasmid, while Clones LT6, LT8, and LT9 were transformed with a linearized form of the same plasmid.

| Clone | Number of reads | Total bases | Mean read length | Mean read quality | Median read length | Median read quality | Read length N50 | Longest read |
| --- | --- | --- | --- | --- | --- | --- | --- | --- |
| Mixed transformants | 151,506 | 1,825,008,054 | 12,045 | 11 | 9,821 | 11 | 15,287 | 83,320 |
| Clone 1 (shallow) | 124,327 | 423,236,197 | 3,404 | 8.8 | 1,547 | 9.4 | 7,662 | 60,490 |
| Clone 5 | 96,988 | 289,944,147 | 2,989 | 8.6 | 1,360 | 9.2 | 6,684 | 59,098 |
| Clone 8 | 83,400 | 307,873,385 | 3,691 | 8.6 | 1,590 | 9.3 | 8,837 | 68,412 |
| Clone 1 (deep) | 708,542 | 7,730,398,077 | 10,910 | 10.9 | 5,733 | 11.8 | 22,989 | 393,346 |
| Clone LT6 | 440,212 | 2,569,220,207 | 5,836 | 12 | 2,739 | 12.1 | 13,490 | 93,794 |
| Clone LT8 | 748,516 | 4,241,519,907 | 5,666 | 11.9 | 4,298 | 12.1 | 9,560 | 154,837 |
| Clone LT9 | 388,980 | 2,353,296,042 | 6,049 | 12.1 | 2,536 | 12.2 | 15,946 | 124,150 |

### Southern hybridization identifies transgenes on high molecular weight species

To gain additional perspective on the fate of transgenes in this first mixed population of transformants, genomic DNA was extracted and run on an agarose gel in three forms: untreated, digested with *Bgl*II, and digested with *Hin*dIII. Both of these 6 base-pair restriction endonucleases cut the plasmid only once. This gel was subsequently used for a Southern blot with a probe against the plasmid-borne neomycin resistance marker gene, neo^R^. This experiment aimed to visualize small molecules (under the ∼30 Kbp resolution limit of a 1% agarose gel) in the undigested DNA sample that could correspond to a stably inherited episomal transgene.

Transgene DNA could be detected in the undigested DNA lane (Figure 3); a single high molecular weight (HMW) band to which the probe hybridizes was seen to migrate higher than the top (20 Kbp) ladder band. In the two genomic DNA digest lanes, a variety of restriction products are apparent, but the Southern hybridization shows very strong signal to a band the same size as the digested plasmid control, with additional weak signal to bands slightly larger and smaller.

**Figure 3.**
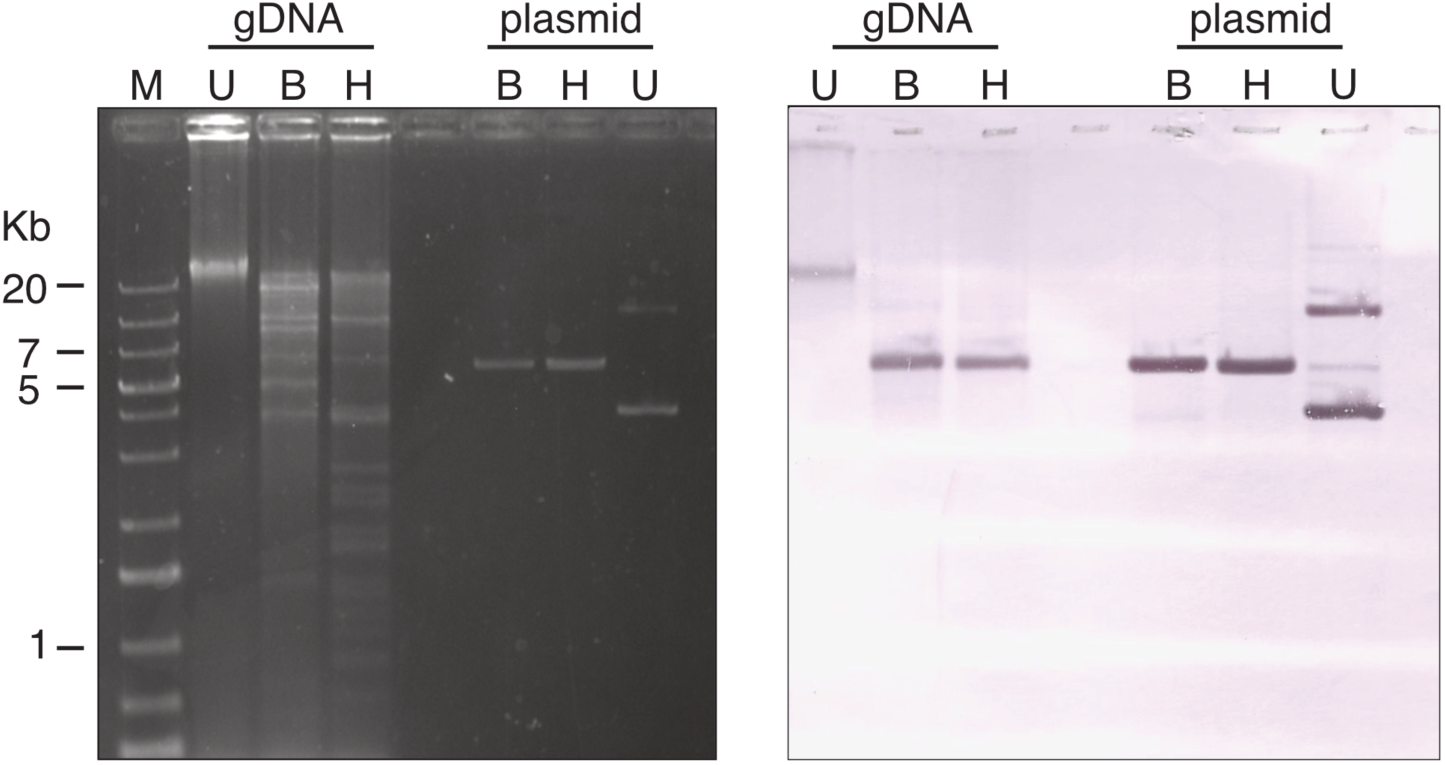
Southern blot detection of the neo^R^ gene in transformed *Acanthamoeba castellanii* genomic DNA. The agarose gel on the right was used for the Southern blot on the right. Genomic DNA (gDNA) from *Acanthamoeba* transformed with pGAPDH-EGFP was run undigested and two aliquots from the same sample were digested with each of two different restriction endonucleases, *Bgl*II (B) and *Hin*dIII (H). These endonucleases each have a 6 base pair recognition site that occurs once within the plasmid. The purified pGAPDH-EGFP plasmid was run as a control, with the same three treatments. The ladder in the leftmost lane is GeneRuler 1 Kb plus DNA ladder, with select band sizes marked alongside. The DNA was transferred to the nylon membrane on the right through a standard alkaline transfer and the neo^R^ probe was hybridized to it. The probe was visualized directly on the membrane with the Thermo Scientific Biotin Chromogenic detection method.

### Nanopore sequencing of monoclonal transformant lines

Three monoclonal transformant isolates were selected for DNA extraction and nanopore sequencing. These three samples were barcoded and sequenced together in a single MinION run (Table 1; ‘Clone 1 shallow’, ‘Clone 5’, ‘Clone 8’). To screen for potential chromosomal integrations of the transgenes, all nanopore reads with hits to the full plasmid sequence were mapped against the wild-type reference genome. Read mapping was done using the soft clipping option, which allowed putative genome-plasmid junctions to be visualized using the Integrative Genomics Viewer^16^ and then verifying that the clipped part of the read was indeed plasmid-derived. One of these monoclonal isolates, ‘Clone 1’, was chosen for repeat DNA extraction and sequencing in the absence of other barcoded samples (Table 1; ‘Clone 1 deep’). The resulting data were analyzed similarly. One putative integration locus on Chromosome 5 was found to be completely spanned by seven nanopore reads, which extend into the unique genomic sequence on both sides. The integrated DNA was found to be a small, 365 bp fragment of the plasmid, corresponding to the protein coding sequence for EGFP. Many of the reads demonstrating putative integrations in these monoclonal cultures contained tandem repeats of the plasmid sequence similar to those described above from previous sequencing experiments. The plasmid repeat units of these arrays were typically all oriented in the same direction, but occasionally there were several repeats in one direction, then a segment of fragmented plasmid sequence, then a few intact repeats in the opposite direction.

### PCR confirmation of a single putative transgene integration

Of the four putative integration loci selected for PCR verification, only one could be confirmed, and only on one end of the integrated plasmid array. This locus was the one covered by seven long nanopore reads that fully spanned the integration. Amplicons from this successful reaction were sent to GeneWiz for Sanger sequencing which confirmed their identity and therefore confirming chromosomal integration of the transgene. For the plasmid-genome junctions on the other end of this locus and both ends of the other three loci, PCR experiments either showed no amplification or amplification of non-specific products, depending on the number of PCR cycles, amount of DNA template, and stringency of the annealing temperature.

### Transformation of *Acanthamoeba castellanii* with linearized plasmid

To test whether rolling-circle replication was involved in the formation of plasmid concatemers, the transformation experiment was repeated with plasmid that had been linearized by the restriction endonuclease *Psi*I, which cuts a single time in the pGAPDH-EGFP plasmid. Monoclonal isolates were derived from the population of successful transformants as described. DNA was extracted from three clones, and all three samples were barcoded and sequenced together in the same nanopore sequencing experiment (Table 1; ‘Clone LT6’, ‘Clone LT8’, ‘Clone LT9’). The number of inferred integrations from the deeply sequenced clone transformed with circular plasmid and the three clones transformed with the linearized plasmid was surprisingly large: 1,251 putative integrations in the clone transformed with circular plasmid, and 280, 187, and 109 inferred integrations in the three clones transformed with linearized plasmid (Table 2).

**Table 2.** Summary of nanopore sequence data bearing on the fate of transgene DNA from four *Acanthamoeba* transformants. Plasmid topology refers to whether the clone was transformed with circular pGAPDH-EGFP or pGAPDH-EGFP linearized by *Psi*I. Apparent integrations are loci where one or more nanopore reads contain genomic sequence from that locus joined to plasmid sequence. The predicted number of artefactually chimeric junctions is determined by calculating the background rate of chimeric reads in a sequencing run, and applying this rate to the number of plasmid containing reads to estimate how many may be chimeric. The reads with one or two telomeres refer to reads where one or more plasmid copies are found in a read with telomeric repeats on one or both ends.

| Clone | Plasmid topology | Estimated genome coverage | Inferred integrations | With 2 reads | With >2 reads | # of artefactual chimeras | 1-telomere reads | 2-telomere reads |
| --- | --- | --- | --- | --- | --- | --- | --- | --- |
| C1 | circular | 100X | 1251 | 25 | 1 | 936 | 288 | 3 |
| LT6 | linear | 50X | 280 | 0 | 0 | 336 | 85 | 3 |
| LT8 | linear | 100X | 187 | 2 | 0 | 438 | 32 | 1 |
| LT9 | linear | 65X | 109 | 1 | 0 | 185 | 45 | 0 |

### Estimating the rate of chimerism in nanopore sequencing experiments

To evaluate how many putative plasmid-genome junctions were the result of read chimerism, the background rate of read chimerism in transformant sequencing datasets was estimated and applied to the total number of plasmid-containing reads. This analysis showed that a strikingly high number of the putative chromosomal integrations of the transgenes could potentially be artefacts generated by read chimerism. Of the 1,251 inferred integrations in Clone 1, read chimerism could be responsible for as many as 936 of them. In Clone LT6 where 280 integrations were inferred, read chimerism was calculated to explain up to 336, in Clone LT8 up to 438 putative integrations could be explained by chimerism while only 187 were inferred, and in Clone LT9, 185 putative integrations could be chimeric, while only 109 were inferred.

To assess the reliability of nanopore sequence-based putative integrations from another angle, putative integration loci that were supported by two or more nanopore reads were identified. Consistent with the read chimerism findings, a low number of putative integrations with more than one read supporting them were found. (Table 2). In Clones 1, LT6, LT8, and LT9 respectively, there were 25, 0, 2, and 1 inferred integrations with 2 or more reads supporting them.

### Illumina sequencing transformed clones reveals extremely high plasmid sequence abundance

Given the uncertainty surrounding nanopore-based evidence for transgene integration, the genomes of two independent *A. castellanii* clones were deeply sequenced using Illumina short read technology. For Clones LT6 and LT9, ∼233.5 million and ∼449.7 million reads were obtained, respectively (30.2 Gbp and 56.2 Gbp of sequence, corresponding to 485X and 910X genomic coverage; Table 3). Two strategies were used to search for putative transgene integrations in these data. First, the set of long reads representing putative integrations were collected into a single file for each clone, and the corresponding Illumina read set was mapped against these putative integrations. The data were then visualized and investigated for short reads mapping across putative plasmid-genome junctions. The second method involved BLAST comparisons of the plasmid, the putative LT6 and LT9 integration reads, and the total set of LT6 and LT9 Illumina reads.

**Table 3.** Illumina and nanopore read coverage depth in Clones LT6 and LT9 when mapped against the wild-type *Acanthamoeba castellanii* Neff genome and against the pGAPDH-EGFP sequence. Coverage values have been rounded to the nearest 5 for genome coverage and to two significant figures for the plasmid coverage for ease of comparison.

| Clone | Genome<br>Nanopore | Plasmid<br>Nanopore | Genome<br>Illumina | Plasmid<br>Illumina |
| --- | --- | --- | --- | --- |
| LT6 | 50X | 12,000X | 485X | 150,000X |
| LT9 | 65X | 18,000X | 910X | 270,000X |

Neither method identified any Illumina reads supporting the putative integrations of plasmid DNA into *Acanthamoeba* chromosomes. To test the hypothesis that the Illumina datasets may be artifactually depleted in plasmid sequence, leading to the observed lack of evidence for integration, the LT6 and LT9 Illumina read sets were mapped onto the sequence of the pGAPDH-EGFP plasmid. The results (Table 3) show that the plasmid was present in extremely high abundance in both LT6 and LT9 cells, albeit in the absence of evidence for how it was being maintained.

### Potential transgene minichromosome in *Acanthamoeba castellanii*

To better understand the ultra-high sequence coverage of plasmid DNA in our nanopore and Illumina sequence data, nanopore reads from all the transformant clones were searched for reads that had one or more plasmid copies flanked on both ends with telomeres, such that they represented a transgene-containing ‘minichromosome’. At least one was found in each of the deeply sequenced transformant clones, ranging from about 30 Kbp to 60 Kbp in length, between 5 and 9 plasmid copies in tandem. Since the actual counts of telomere-flanked reads were very low, all plasmid arrays with telomeres on at least one end were identified to account for the possibility that most reads were not long enough to capture both telomeres. The tabulated counts of these reads for each clone are presented in Table 2.

We next sought to visualize the hypothesized transgene minichromosome using gel electrophoresis and Southern hybridization. In this experiment, rather than using two different restriction enzyme digests, we aimed to degrade away as much wild-type genomic sequence as possible without cutting plasmid arrays. For this, *Sac*I, *Nhe*I, and *Not*I were chosen due to each one having at least a few thousand cut sites in the wild-type Neff genome, and none in the pGAPDH-EGFP sequence. More specifically, *Nhe*I cuts a predicted 2,565 times in the 35 chromosome-scale scaffolds of the wild-type Neff genome^11^, for an expected average of roughly 17 Kbp between cut sites, while *Not*I cuts a predicted 3,790 times in those scaffolds for an expected average of roughly 11 Kbp between cuts, and *Sac*I cuts a predicted 24,049 times in those 35 scaffolds for an expected average of roughly 2 Kbp between cuts. A 0.5% agarose gel (for improved resolution) was run with wild-type Neff gDNA, LT9 gDNA, and pGAPDH-EGFP. Each sample was run in undigested form as well as digested with the cocktail of restriction enzymes described above. A Southern blot was again performed with a probe against the neo^R^ gene from the plasmid.

From the gel (Figure 4), the WT and LT9 digested lanes appear to have approximately the same banding pattern. Interestingly, in the LT9 undigested gDNA lane there is a distinct extra band appearing just below the combined HMW DNA band. Given the expectation for the transformants to have a 30 to 60 Kbp molecule bearing the transgenes, this distinct extra band appears to be a candidate, but the Southern blot shows no hybridization of the neo^R^ probe to this extra band, instead hybridizing to the high molecular weight DNA band. This extra band thus appears not to be the hypothesized linear episome, but its identity is not clear and could not be narrowed down by revisiting the sequence data from this clone, nor from PFGE of the undigested wild-type and LT9 samples (data not shown).

**Figure 4.**
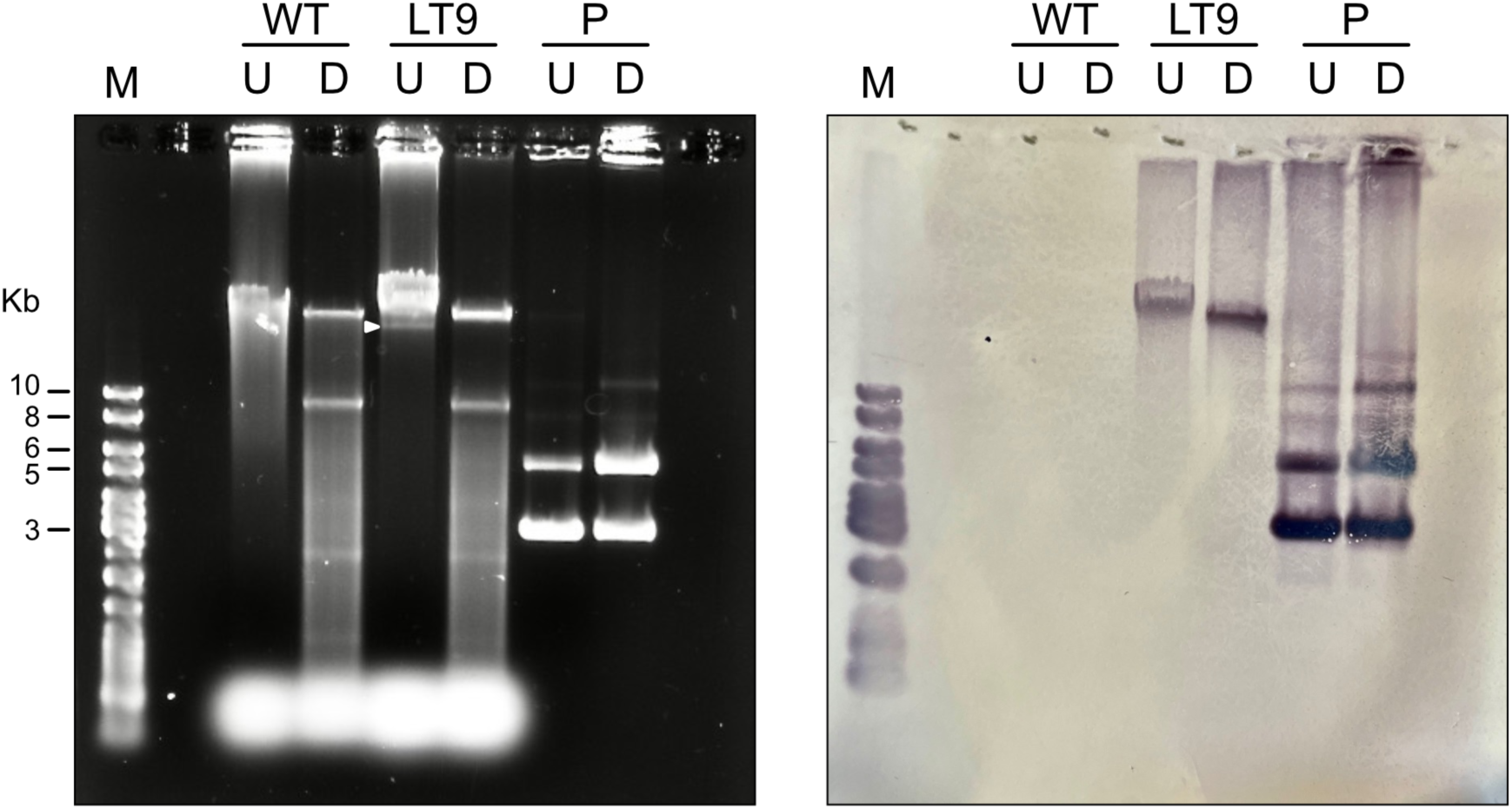
A Southern blot to detect extrachromosomal transgenes in transformed *Acanthamoeba castellanii* genomic DNA. The agarose gel (left) was used for a Southern blot / hybridization (right). Genomic DNA from wild-type (WT) *A. castellanii* strain Neff and Clone LT9 was run undigested (U) and digested with *Sac*I, *Nhe*I, and *Not*I (D). These endonucleases cut frequently within the wild-type genome and do not cut within the plasmid. The purified pGAPDH-EGFP plasmid (P) was run as a control, with the same treatments. The ladder in the leftmost lane is GeneRuler 1 Kb plus DNA ladder, with select band sizes marked alongside. An arrowhead marks a band that appears in undigested LT9 but not wild-type. The DNA was transferred to the nylon membrane on the right through a standard alkaline transfer and the neo^R^ probe was hybridized to it. The probe was visualized directly on the membrane with the Thermo Scientific Biotin Chromogenic detection method.

## Discussion

The goal of this study was to determine the fate of transgenes in *Acanthamoeba castellanii* after artificial transformation experiments with circular and linearized forms of the same plasmid. Nanopore and Illumina sequencing, PCR, Southern blotting, and PFGE were all used to determine whether the transgenes were episomal or chromosomally integrated, and if any modifications to them had taken place. The results are complex, but on balance favour the episomal maintenance of the transgenes in *Acanthamoeba* on a linear molecule that contains a tandem array of plasmid sequence flanked by telomeric repeats.

Our results unambiguously indicate that *Acanthamoeba* maintains the transgenes in tandem arrays of between 5 and 10 plasmid copies on average. It is unclear whether this size range would consistently apply to any future transformation experiments on this organism with varying parameters, nor to natural acquisition of foreign DNA in either circular or linear form. Distinguishing between chromosomal or episomal maintenance of transgenes was not possible using the array of experimental approaches employed. Using the size of the transgene-containing DNA species as a diagnostic criterion was surprisingly inconclusive, even using standard DNA sizing methods like agarose gel electrophoresis. The size of a putative episome, inferred to be in the range of 30 Kbp to 60 Kbp based on sequence data, is much different than the size of most of the nuclear chromosomes in *Acanthamoeba*^11^, although a couple of chromosomes at the lower end of the size range are around 100 Kbp. However, 30-60 Kbp is still above what can easily be resolved on a standard agarose gel. Therefore, while Southern blot experiments placed the transgenes in the same compressed, HMW band on as the genomic DNA, the episome may not be expected to resolve as a separate band on such gels. Peng, Omaruddin, and Bateman^5^ arrived at a similar conclusion in their original description of the transformation protocol for *Acanthamoeba*, which used the circular form of the plasmid. Using Southern hybridization, they concluded that the transgenes must be on a molecule of at least 12 Kbp in size and present in several copies, but could not differentiate between chromosomal integration, the formation of a linear episome, or a circular molecule composed of multiple plasmid concatemers. Use of PFGE to try and resolve an episomal band from intact chromosomal DNA provided no additional insight; no clear band in the size range that would be expected based on the sequence data alone could be observed (data not shown). A future experiment using a PFGE-resolved gel as substrate for Southern blots with probes against telomeres and the neo^R^ resistance marker may clarify the situation.

Overall, sequence-based analysis arguably shifts the balance of probabilities in favour of episomal maintenance of transgenes in *Acanthamoeba* rather than strictly chromosomal integration, for several reasons. The first is that the evidence for the latter is unconvincing. The discrepancy between the read support of any given putative integration and the overall genome coverage demands explanation. Depending on the specific integration and the transformed strain analyzed, this could be a 25- to 100-fold difference; while putative plasmid integrations were supported by only one or two reads, the genome coverage of the clones ranged from 50X to 100X. The explanation held throughout much of this investigation was that the suspected polyploidy of *Acanthamoeba*, estimated to be ∼25*n* by Byers^18^, could account for this discrepancy if an integration took place in only one copy of a chromosome. However, finding a single supporting read for an integration in a sequence dataset of 100X genome coverage makes even this scenario unlikely.

This explanation (i.e., transgene integration into one or a few chromosomal copies) is further weakened by the ultra-high abundance of plasmid sequence in both Illumina and nanopore sequence data: the coverage of a single copy of the plasmid was about 250 to 300 times higher than the genome coverage in clones LT6 and LT9 (Table 3). Even giving the integration hypothesis maximum ‘benefit of the doubt’ does not account for this finding; assuming each of the 280 putative integrations in LT6 had an array of 11 plasmid units (the longest array found in any of the data sets), this explains 3,080 total copies of the basic plasmid unit, which is only 62 times higher than the overall genome coverage in the corresponding read set, falling short of the observed 250 to 300X coverage.

Consideration of read chimerism further weakens this hypothesis. Based on the background rate of chimeric reads estimated in the LT6 nanopore sequencing run, chimerism could explain the appearance of up to 336 reads corresponding to artefactual junctions between plasmid and genomic sequence, which exceeds the observed number. It should be noted that the criteria used to identify chimeric reads were relatively stringent; one could propose situations of true chimerism that would be rejected by the criteria implemented. This means that the estimated rate of chimeric reads, if inaccurate, would likely be an underestimate.

The presence of telomere-flanked plasmid arrays in the nanopore reads obtained from multiple transformed clones provides evidence for the existence of a transgene that could be replicated and maintained as an episome. This hypothesis shares some of the weaknesses of the integration hypothesis, but to a lesser extent. Our finding of three or fewer reads in each clone where a plasmid array is flanked by telomeres on both ends is difficult to explain. Even when accounting for reads with telomeric repeats on only one end, the counts are quite low compared to total plasmid sequence abundance. However, the issue of read chimerism is less problematic for this hypothesis. For chimerism to account for all plasmid-telomere junctions observed in each clone, each chromosome would be expected to have telomeres in excess of 100 Kbp on each end. This is based on using the estimated rate of chimerism in each clone to extrapolate how much total telomeric sequence would be needed for all observed plasmid-telomere junctions to be chimeric, then dividing this by the number of chromosome ends. For comparison, mammalian telomeres are typically from 10 to 30 Kbp, and yeast telomeres are only a few hundred base pairs in length. This would also make each of *Acanthamoeba*’s telomeres at least the same length as the total predicted non-telomeric sequence of chromosome 35. It is thus more difficult to reject the existence of a linear episome composed of plasmid concatemers and flanked by telomeres than it is to reject most of the putative chromosomal integrations inferred from the sequence data.

All this said, there is a single chromosomal integration of transforming DNA in Clone 1 that clearly cannot be rejected. This is the only case where long reads fully span a putative integration site, and is also supported by several times more reads than the others. It is also the only example where molecular support could be obtained (i.e., PCR). It is thus highly likely that the putative transgene integration into *A. castellanii* chromosome 5 of transformant line ‘Clone 1’ is genuine. However, this instance is also unlike any of the other plasmid sequences found in the four analyzed clonal lines in that only a 365-bp fragment of the plasmid is integrated. This sequence comes from the EGFP gene and does not contain either of the termini required for proper expression, so it seems unlikely to have any biological influence on the cell, including not contributing to resistance against the selecting antibiotic in culture. Extrachromosomal copies of the transforming plasmid thus must be responsible for antibiotic resistance in this clone, as is expected in the others. This integrated DNA would also not have been detected by the probe against the neo^R^ gene, but this clone was not probed experimentally anyway.

A strikingly similar situation was observed in the fungal pathogen *Histoplasma capsulatum* by Woods and Goldman (1992^19^ and 1993^20^). Transforming plasmids in *H. capsulatum* were either chromosomally integrated, or modified into a linear plasmid closely resembling the ones observed here in *Acanthamoeba*, including addition of telomeric repeats followed by maintenance of the episome at high copy number. Both of these potential destinations for the transgene DNA showed tandem amplification of the transforming plasmid. Transformation with circular or linear plasmid does not appear to influence transgene fate. Although only one chromosomal integration was observed here for *Acanthamoeba*, our overall findings are nearly identical to those in *Histoplasma*^19, 20^.

Similar outcomes have been described in the fungal genetics literature, and in other protists. Examples include the fungi *Cryptococcus neoformans*^21,22^ and *Fusarium oxysporum*^23^ and the ciliate *Paramecium tetraurelia*^24^. Of these examples, none exhibit tandem duplication of the transforming plasmid as clearly as *Acanthamoeba* or *Histoplasma*, with *F. oxysporum* potentially having partial duplication and rearrangement of the transforming DNA while the transforming plasmid remains unit-sized in the other two examples. All three do appear to add telomeric repeats to the transforming DNA, apparently without any particular motif required for template the addition. All these organisms must either have sufficiently permissive replication systems such that no specific replication origin is needed or the telomeres themselves can serve as a replication origin; the literature suggests that one or the other of these options can be true depending on the lineage.

Among the fungal systems discussed here, all three exhibited the ability to maintain transforming DNA through both chromosomal integration and the formation of autonomous linear episomes. In each of the three, experiments demonstrate that transforming DNA can be maintained through both means simultaneously. These findings support the inference that *Acanthamoeba* ‘Clone 1’ in our study has generated a linear episome from the transforming DNA while also containing one bona fide chromosomal integration. Whether this integration occurred independently of linear episome formation or whether a linear episome served as a substrate for this integration is unknown.

With respect to the mechanism responsible for generating tandem arrays of the plasmid sequence, evidence indicates that the transforming plasmid was concertedly duplicated in some way. It is unlikely that random ligation of several plasmids gave rise to these arrays, based on the non-random orientation of the plasmid units within a given array. Intriguingly, the telomeres do not appear to join the plasmid array at the position where the original transforming plasmid was linearized. The nucleotide position on the plasmid where telomeres were joined was used as a proxy to estimate the diversity of episomes within each transformed clone. In each respective clone, there appears to be one dominant form of the episome, with a frequency of 20% - 50% depending on the clone, followed by one to a few minor forms with frequencies between 5% and 20%, and the remainder of the observed episome-like reads are effectively ‘singletons’ in our data. This would suggest each clone has a few different versions of the episome that are replicated somewhat more efficiently than the rest, with a large diversity of rare versions. It is unclear whether this diversity of forms arose from a single episome due to instability, or whether multiple episomes formed at the time of transformation due to the large quantity of transforming DNA, with some being held at higher copy number than others. It seems plausible that the latter scenario is true and the difference in frequency of the different forms may be caused by differences in replication efficiency due to unknown factors.

At least two hypotheses can explain why there are so many observed positions where telomeres join the plasmid. Perhaps the instability known to be intrinsic to tandemly duplicated sequences caused sufficient rearrangement to create this variation in plasmid-telomere junction sites. Examination of the episome-representing reads provides support for this hypothesis; the arrays are often composed of neatly duplicated plasmid copies but there are cases where there is significant fragmentation of the plasmid units within these episomes, presumably due to recombination. Another possible explanation is that degradation of the ends of arrays, whether passively or by enzyme-mediated processes, caused the ends to vary until telomeres were added to protect the plasmid arrays and preserve the structure that existed at that moment in time. These two possibilities are not mutually exclusive; both could contribute to the observed variation in plasmid-telomere junctions in our *Acanthamoeba* transformants.

If the hypothesis favoured here is true — i.e., that *Acanthamoeba* maintains transformed DNA as an autonomous linear plasmid — this would have evolutionary implications as well as implications for molecular biology experimentation in this system. Here we must distinguish between *autonomy* and *stability* of these extrachromosomal elements. Autonomously replicating plasmids can be recognized and propagated by the endogenous DNA replication machinery, but this does not guarantee their stability (i.e., longevity) across generations. In non-integrated elements, selection must typically be maintained to prevent loss of the plasmid over time.

Extrachromosomal elements such as the ones described here for *Acanthamoeba* are employed as a molecular biology tool by fungal geneticists, who append telomeres to a variety of genetic constructs and have the cell maintain them autonomously. In fungi, autonomously replicating plasmids provide great utility for genetic experiments by virtue of high transformation efficiency and high copy number; transforming DNA that cannot replicate autonomously has to be chromosomally integrated which is comparatively much less efficient. Some yeasts can maintain circular plasmids autonomously provided they have an ARS, but in many of these fungi, the addition of telomeres to linear plasmids creates highly efficient genetic constructs. This knowledge could be used to develop genetic tools for highly efficient transformation in *Acanthamoeba* to study its biology more intensively. The same concept could possibly be applied to other nascent protist model systems where transformation is difficult or not yet achieved.

From an evolutionary perspective, our findings suggest a broader distribution of episomal transgene maintenance across eukaryotes than previously thought. With the exception of the alveolate ciliate *Paramecium tetraurelia*, the formation of telomere-bearing linear episomes has only been observed in fungi. While Amoebozoa is the closest supergroup to Obazoa, which contains fungi, this should still prompt molecular biologists working on protists from other parts of the eukaryote tree to be alert to similar phenomena. The observation of this type of molecular biology in the two major groups of Amorphea (i.e., Amoebozoa plus Obazoa), as well as in an alveolate, thought to be on the opposite side of the eukaryote root^25,26^, could indicate that it is widespread among eukaryotes.

A second evolutionary implication of episomal transgene maintenance in *Acanthamoeba* is the role it may play in fine-scale genome evolution. Many transformation protocols serve to facilitate access of the transforming DNA to the nucleus, but do not themselves provide an obvious mechanism by which the DNA can be maintained once it has arrived there. There are exceptions, such as in *Agrobacterium*-mediated transformation, where the bacterium encodes machinery for integrating the DNA^27^, or transformations paired with gene editing systems like CRISPR-Cas9. However, in presenting exogenous DNA to the nucleus, most transformation protocols may not differ from this point onward from exogenous DNA acquired in a natural environment (assuming it can access the nucleus), which may serve as substrate for lateral gene transfer (LGT). The spontaneous generation of autonomously replicating plasmids described in this study could allow exogenous DNA to persist longer in the nucleus, which would greatly expand the window of opportunity for other requisite steps of LGT, such as chromosomal integration and gene expression, to occur.

One can even imagine some of the steps involved in expressing the newly acquired genes to take place prior to integration; in fact, the unstable and recombinogenic nature of the tandemly duplicated transforming DNA in this study could facilitate the acquisition or invention of promoters and other necessary elements, if tandem duplication also happens to naturally acquired DNA. Aside from providing a standing source of foreign DNA for integration, autonomous plasmids could be a substrate for innovation. They may even begin expressing genes that provide a selective advantage to the recipient cell *before* chromosomal integration, encouraging maintenance of the new genetic material until it can be incorporated more permanently. This hypothetical model could help tip the scales in what is sometimes thought to be a highly improbable series of events.

## Conclusions

We have shown how long-read, single molecule sequencing can serve as a high throughput readout for molecular biological experimentation. While still not trivial, a wide range of experimental possibilities can be explored with genome-wide sequence read sets much more quickly than with traditional molecular biology experiments. However, it also leaves room for ambiguities that must be explored and resolved through experimentation.

Moving from methodology to biology, our results show that *Acanthamoeba* appears to duplicate circular and linear plasmid DNA into a tandem array and add telomeres to the ends such that it can be autonomously replicated in the nucleus. This genetic capability has not been observed in *Acanthamoeba* before, and would suggest that the organism’s telomerase is capable of acting on substrates without existing telomeric repeats, and that its replication machinery is able to replicate non-chromosomal DNA, potentially by recognizing the newly added telomeres. We cannot rule out the possibility of chromosomally integrating exogenous DNA; indeed, our data suggest that *Acanthamoeba* has the genetic flexibility to use autonomously replicating elements and chromosomal integration to retain foreign DNA, possibly simultaneously. These observations can be exploited by molecular biologists to improve the efficiency and flexibility of genetic manipulation in this organism. At the same time, these capabilities hint at the existence of a molecular gateway for lateral gene transfer in nature, as has been suggested to occur in *Acanthamoeba* and other Amoebozoa based on the presence of bacterial and viral genes in their genome^11,28–30^. The formation and maintenance of linear episomes from foreign DNA could extend the window for integration to occur.

Our findings also contribute to a broader evolutionary discussion on the ubiquity of such genetic mechanisms and their role in genome evolution across eukaryotes. The distribution of similar processes across major eukaryote groups could reflect a widespread or even universal mechanism for sampling extrachromosomal DNA in the nucleus and potentially retain it more permanently. Such a system could be useful for retaining endogenous DNA that has been expelled from its chromosomal home and would otherwise no longer be replicated. This could also serve as a mechanism for facilitating eukaryote-eukaryote LGT across the eukaryotic tree via the creation, maintenance and transfer of episomes. Beyond virus-mediated transfer, the mechanisms underpinning LGT in eukaryotes is largely a black box^31^ and elucidation of previously unrecognized genetic capabilities such as episomal maintenance could go a long way toward explaining how LGT contributes to the evolution, diversification, and ecological adaptation of eukaryotic cells in nature.

## Acknowledgements

This research was supported by a grant from the Gordon and Betty Moore Foundation (GBMF5782). M.J.C. was supported by graduate student scholarships from the Natural Sciences and Engineering Research Council of Canada and Dalhousie University.

